# Antigen binding kinetics are quite different for B-cell receptors and free antibodies

**DOI:** 10.1101/2022.07.19.500451

**Authors:** Miguel García-Sánchez, Mario Castro, José Faro

**Author notes:** All the authors contributed equally except MGS that also performed all the stochastic simulations.

## Abstract

Since the pioneering works of Berg and Purcell, discriminating between diffusion followed by binding has played a central role in understanding cell signaling. B-cell receptors (BCR) and antibodies (Ab) challenge that simplified view as binding to antigen follows after a chain of diffusion and rotations, including whole molecule rotation, and independent tilts and twists of their Fab arms due to their Y-shaped structure and flexibility. In this paper, we combine analytical calculations with Brownian simulations to derive the first-passage times due to these three rotations positioning the Fab paratopes at a proper distance and orientation required for antigen binding. Applying these estimations and those for 2-dimensional (2D) and 3D translational diffusion of, respectively, BCRs and Abs, we evidence that measuring Ab-Ag effective kinetic binding rates using experimental methods in which the analyte is in solution gives values proportional to the intrinsic binding rates, *k*^+^ and *k*^−^, only for values of *k*^+^ up to 10^9^ s^−1^, beyond which a plateau of the effective 3D on rate between 10^8^ M^−1^s^−1^ and 10^9^ M^−1^s^−1^ is attained. Moreover, for BCR-Ag interactions, the effective 2D on and off binding rates can be inferred from the corresponding effective 3D on and off rates only for values of effective 3D on rates lower than 10^6^ M^−1^s^−1^. This is highly relevant when one seeks to relate BCR-antigen binding strength and B cell response, particularly during germinal center reactions. Thus, there is an urgent need to revisit our current understanding of the BCR-antigen kinetic rates in germinal centers using state-of-the-art experimental assays for BCR-Ag interactions.

**Significance Statement:** In germinal centers, binding between BCRs and antigen (Ag) tethered on the membrane of follicular dendritic cells occurs via two-dimensional (2D) membrane-to-membrane interactions. In contrast, in *in vitro* assays antibody (Ab)-antigen interactions occur with one component in solution. Structurally, there are large qualitative and quantitative differences between BCR-Ag 2D and Ab-Ag 3D translational and rotational diffusion processes, with the 2D translational diffusion being about 1000-fold lower than the 3D one. Moreover, the effective binding kinetics of both BCR-Ag and Ab-Ag interactions strongly deviate from the intrinsic molecular on and off rates. Here we expose this mismatch and, performing numerical and analytical calculations, quantify the ranges for which the experimental in-vitro data is informative on the BCR-Ag binding strength.

---

**O**ur current understanding of antibody (Ab)-antigen (Ag) interactions rests on decades of experiments on diffusion-limited reactions. Antigen binding to B-cell receptors (BCRs) is the first step of Ag-specific activation of B lymphocytes, ultimately leading to their differentiation into Ab-secreting plasma cells. However, although both BCRs and Abs are immunoglobulins (Ig), they critically differ in that while BCRs are membrane Igs possessing a C-terminal tail that spans the B cell membrane until the cytoplasm, Abs are soluble, Ig isoforms that lack the membrane-anchoring and transmembrane tail (1). The basic structure of Igs includes two identical Ag-binding arms or fragments (Fab) that are linked to a single constant arm (Fc) through an aminoacid sequence known as hinge region (Figure 1a) forming their familiar bivalent Y shape (1). Fab arms bind Ags through a small area located at each free Fab end named paratope (*P*), and the particular Ag interface area participating in a Fab-Ag bond is known as epitope (*E*) (Figure 1a). The hinge region linking Fab arms to the Fc arm has a variable length and flexibility specific to each Ig isotype (2). This allows the two Fab arms to pivot independently of each other about the Fc region.

**Fig. 1.**
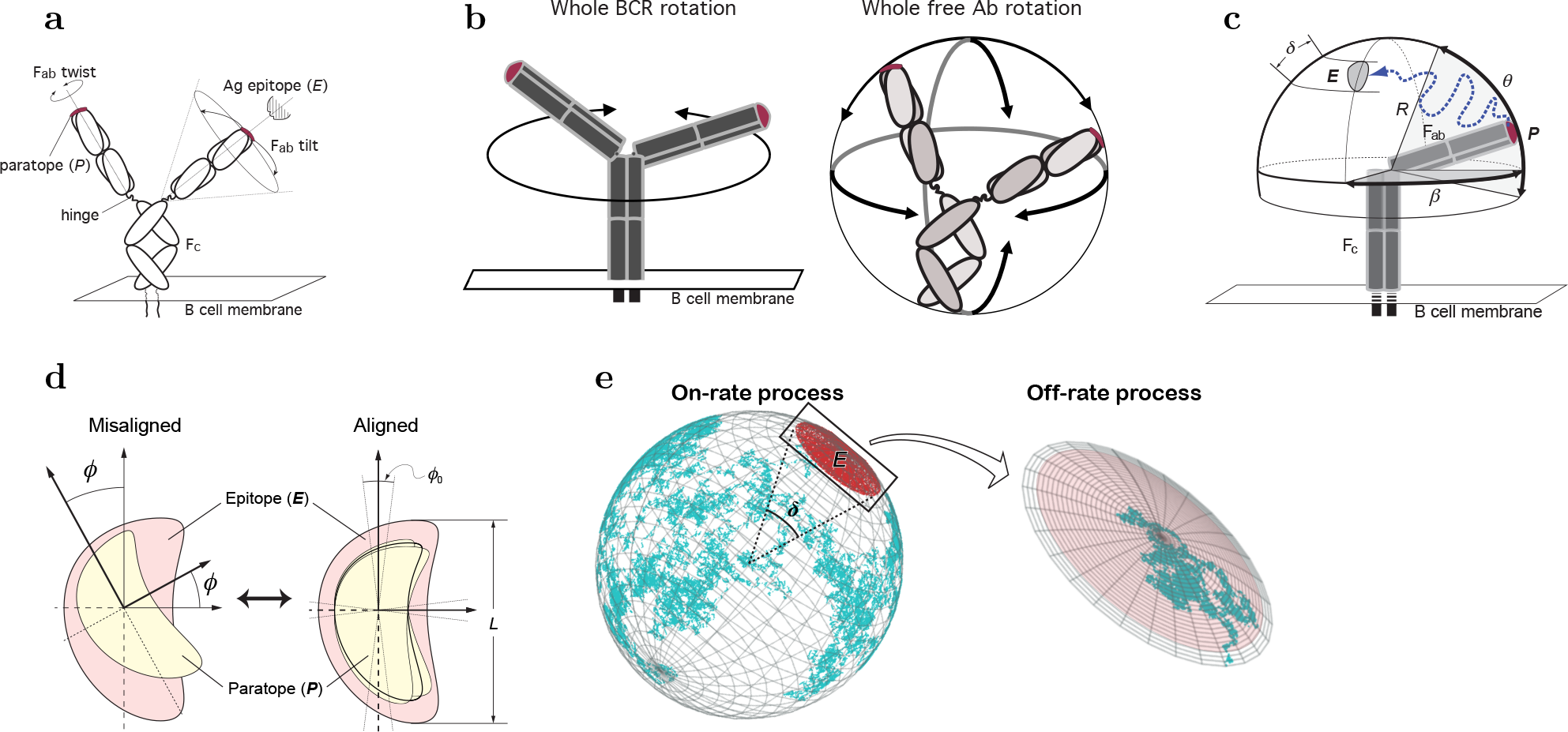
Possible Ig rotations impacting on the position of a paratope *P* relative to an Ag epitope *E*. **a**. The position of each paratope in an Ig (whether membrane-immunoglobulin, mIg, or free antibody, Ab) is independently submitted to two possible Fab rotations, Fab tilt (wagging and wobbling; depicted on the right side), and Fab twist in which each Fab arm rotates along its major axis (from the hinge region to *P*; depicted on the left side). The speed of these two Fab rotations might be different in BCRs and free Abs. **b**. The position of each paratope is also affected by the rotation of the whole Ig. The left side depicts BCR whole rotation, mostly restricted to a rotation along the Fc axis. The right side depicts free Ab whole rotation, which is unrestricted in the three spatial dimensions. **c**, Sketch of the diffusion of *P* due to Fab tilt, assuming the epitope is in a fixed position; this movement is constrained to a spherical cap on a sphere of radius *R* = length of a Fab arm ≈ 84 Å (12) and to Fab altitude and azimuth angles, respectively, *θ* (measured from the *south pole* of the sphere) and *β* (the azimuthal angle of the Fab measured from the location of the epitope). Depicted is a possible path followed by paratope *P* toward epitope *E* (dotted line). The angle *δ* corresponds to an arc on the spherical surface equal to the major length of *E*. For clarity, the outlined mIg is depicted with only one Fab arm. **d**, Alignment or misalignment of *P* and *E* due to Fab twist. *P* rotates with variable angle *ϕ* relative to *E* (left side). Such rotation will align or misalign *P* and *E*, allowing or impeding their chemical binding. For aligned *P* and *E* there is a tolerance angle, *ϕ*_0_, within which the chemical reaction can proceed (right side). The view in panel **d** is along the major Fab axis. **e**, Random Walk view of Igs’ paratope diffusion. During the on-rate process, the center of mass of the paratope diffuses on the sphere (cyan broken line) until finding the epitope (red area); *δ*, the angle of the arc with a length equal to the major length of the epitope. During the off-rate process, the paratope’s center of mass diffuses within the epitope area (pale red) until it escapes (grey area).

During a humoral immune response to protein Ags, B cells interact through their BCRs with their cognate Ag held in immunocomplexes on the membrane of follicular dendritic cells (FDC) (3–5). This receptor-ligand interaction is similar to that of T-cell receptors (TCR) and their ligands in that it takes place in the context of a membrane-membrane interaction, and therefore in 2-dimensional (2D) conditions (6, 7). Yet, the effective kinetic rate constants and affinity constants of BCR-Ag and Ab-Ag interactions in 2D have not been measured until now, except for a couple of Abs (8, 9), due to a lack of adequate, high throughput techniques that are fast and easily implemented in a lab. In contrast, for 3D Ab-Ag interactions. Several available techniques, particularly the so-called Surface Plasmon Resonance (SPR) assay, are widely used to measure the Ab-Ag effective kinetic rate and affinity constants. Traditionally, it has been implicitly assumed that 3D Ab-Ag interactions are a good proxy for 2D BCR-Ag interactions. However, for TCRs this approach has been proven misleading (6).

We have shown before that, in general, protein-protein interactions can be described in detail by considering the three main sequential processes taking place in that interaction, namely: translational diffusion, rotational diffusion, and molecular binding (6). In the case of BCR-Ag interactions (2D conditions) and Ab-Ag interactions (3D conditions), the translational *on* and *off* kinetic rates scale similarly to each other from 2D to 3D conditions (7). However, it is presently unknown whether the corresponding rotational *on* and *off* kinetic rates also scale similarly to each other from 2D to 3D conditions, as well as what are the limits to the effective 2D kinetic rates sensed by B cells and, most important, if and under what conditions the 2D BCR-Ag effective kinetic rates can be inferred from the 3D Ab-Ag effective kinetic rates.

Here we perform a detailed analysis of BCR and Ab paratopes’ rotational rates in 2D and 3D conditions, respectively, using the theory of stochastic narrow escape (10), and first passage processes (11). We combine existing experimental data with analytical calculations and Brownian simulations to provide a complete quantitative picture of the first-passage times associated with the BCR and Ab rotational degrees of freedom. Our results allowed us to estimate the effective *on* and *off* rotational rates, and hence the effective 2D (BCR-Ag) and 3D (Ab-Ag) effective *k*^+^ and *k*^−^ rate constants as a function of the intrinsic or molecular *k*^+^ binding rate. Moreover, we could derive a formula relating the 2D effective kinetic rates to the corresponding 3D effective kinetic rates. Contrary to the traditional implicit assumption, the kinetics of BCR-Ag interactions cannot be inferred, in general, from those of Ab-Ag interactions above a certain *observability threshold*, close to 10^5^ M^−1^ s^−1^ for the 2D effective *k*^+^ rate and 10^6^ s^−1^ for the 2D effective *k*^−^. Consequently, the kinetics of BCR-Ag interactions cannot be inferred, in general, from those of 3D Ab-Ag interactions.

## A first-passage-time formalism of the rotational diffusion steps during BCR and antibody binding to antigen

### The three-step description of kinetic binding

In general, the kinetics of reversible protein-protein interactions consist of three generic processes (mostly occurring sequentially): translation, rotation, and binding. Translational diffusion comprises the spatial approximation of *R* and *L* molecules to a distance that allows binding; we denote this *close-enough* by *RL*\*. In the second step, rotational diffusion, the two close molecules must attain a proper orientation to bind to each other; we denote this close and oriented state as *RL* (in the case of Igs, a paratope has to be properly oriented for binding its epitope). Once properly oriented, they can form a bound complex *C* (6). Equation (1) summarizes these series of states and processes,

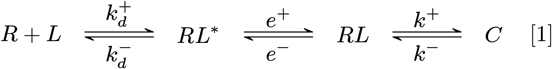

where 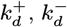 are, respectively, translational diffusion *on* and *off* rate constants, *e*^+^ and *e*^−^ are diffusion *on* and *off* rate constants for effective or global rotation, and *k*^+^ and *k*^−^ are the intrinsic or molecular *on* and *off* rate constants. It is worth noting that De Lisi (13) already emphasized the relevance of this three-step analysis generalizing the diffusion only plus binding framework pioneered by Berg and Purcell (14).

Traditionally, binding has been analyzed experimentally, lumping together those three steps assuming they occur in a single step. The *effective* kinetic rates for this simplified view can be formally derived by assuming a quasi-steady state approximation for the intermediate states [*RL*\*] and [*RL*], which leads to the following expressions for 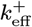 and 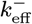 (6):

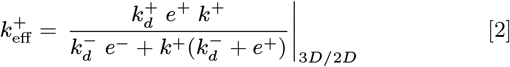

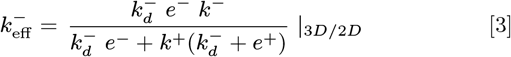

The subscripts 3*D*/2*D* denote that the translational and global rotational diffusion rate constants correspond to either 3D or 2D reactions. Note that *k*^+^ and *k*^−^ are independent of the dimensional condition and, thus, are identical in 3D and 2D interactions. The divisor in these equations determines an *observability threshold* for *k*^+^,

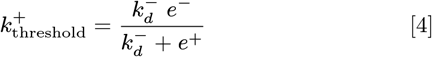

such that for 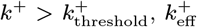 approaches asymptotically a plateau given by 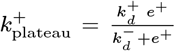, and 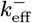 becomes proportional to the dissociation affinity constant (7).

This three-step process can be further analyzed to capture the subtle —yet crucial— role of the Y structure of BCRs and Abs. Due to thermal fluctuations, proteins diffuse randomly, driven by molecular collisions or vibrations, until that motion stops by reaching a binding site. In particular, the wagging and wobbling movements of Ig Fab arms relative to the Fc arm, made possible by the hinge region of the Ig heavy chains, determine the position of the corresponding paratopes relative to the Fc arm and are referred to here as tilt rotation of paratopes or Fab tilt (Figures 1a,c) (2, 15, 16). The *on* and *off* rate constants characterizing this rotation are denoted here 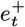 and 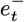, respectively. In addition, a Fab arm can twist about its major axis thanks again to the relative flexibility of the hinge segment (17, 18). This Fab twisting contributes essentially to the orientation or alignment of a paratope relative to an epitope on a relatively immobile Ag molecule tethered to an FDC membrane. We refer to this rotation as alignment rotation (Figures 1a,d), and the *on* and *off* rate constants characterizing this rotation are denoted, respectively, 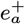 and 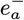. Last, the possible rotation of the whole BCR about the Fc axis, like a *spinning top*, or the rotation of an Ab in solution as a whole (Figure 1b) contributes to the global paratope rotation with *on* and *off* rate constants denoted 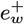 and 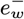, respectively. Due to such a complex combination of different rotational diffusion modes affecting the relative position of paratopes of BCRs and free Abs, the above-mentioned global rate constants *e*^+^ and *e*^−^ are, in fact, compound quantities of the more elemental rates characterizing the whole molecule, Fab tilt and Fab alignment rotations.

The global *on* rotation takes place in two steps. First, a Fab paratope *P* becomes close enough to an epitope *E* through whole Ig and Fab tilt rotation, and then *P* and *E* become correctly oriented by a twist or alignment rotation. The first step, the close approaching of *P* and *E*, combines the whole molecule and Fab tilt rotation processes taking place in parallel or in any temporal order. Hence its rate is the sum of the two individual processes, 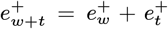, and thus the corresponding mean time is given by 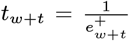. However, the *on* alignment rotation follows *sequentially* after the approaching of *P* and *E*, hence the mean rotation time of the two steps (approaching and alignment) is the sum of their respective mean times, that is, *t*_*on*_ = *t*_*w*+*t*_ + *t*_*a*_ and, consequently, 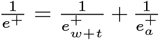. At variance with *e*^+^, the global *off* rotation rate, *e*^−^ (*i.e*., from the *RL* state to the *RL*\* state), is determined by the *whole* and *tilt* off rotations, which can take place in *parallel* or in any temporal order. This can be described in terms of the following reaction scheme:

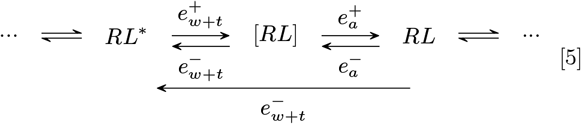

And according to the above:

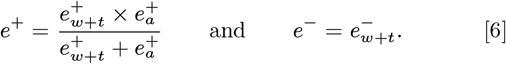

### 2D compound tilt+whole molecule rotational rates of BCRs

In BCRs, the anchoring to the cell membrane has two main effects related to the degrees of freedom of paratope movements. First, its whole rotation motion is restricted to essentially the axis normal to the cell membrane in a highly crowded molecular environment. Although the exact value of the whole BCR rotational diffusion coefficient, *D*_*w*_(2*D*), has not been experimentally determined, one can take as a reference value that of the epidermal growth factor receptor (19), which is of the order 10^3^ rad^2^/*s*. This is three orders of magnitude smaller than the 2D tilt rotational diffusion coefficient *D*_*t*_(2*D*) (Table S3). Second, the Fab tilt flexibility determines the spherical surface domain Ω that the paratopes cover (Fig. S5). The different Ig isotypes and subclasses in humans and mice have different hinge regions and, consequently, they differ substantially in flexibility and tilt rotation. The flexibility in the polar angle *θ*, i.e, the flexibility of the Fab arms with respect to the Fc region (Fig. 1c), has been characterized (SI, section A and Table 2).

**Table 1.**
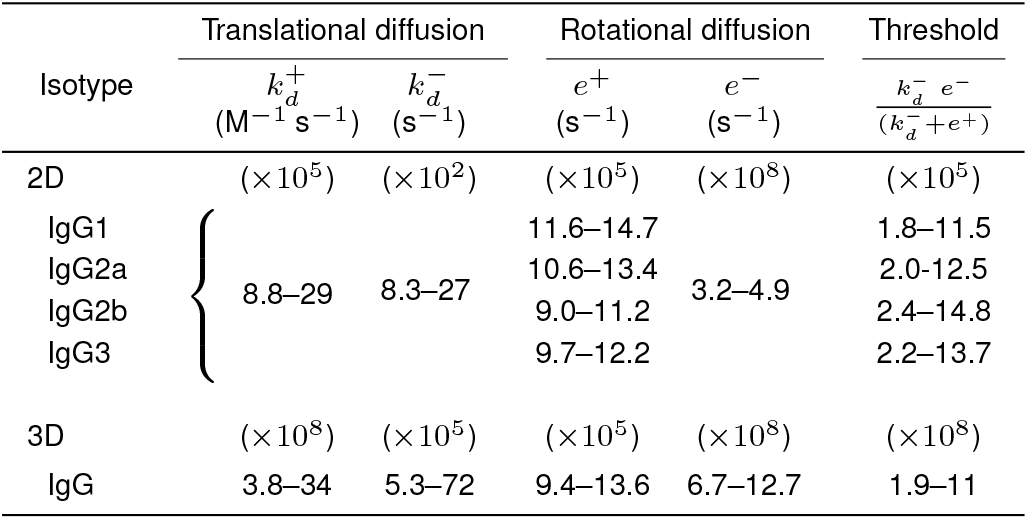
Estimated ranges of translational and rotational diffusion rate constants for Igs in 2D and 3D conditions, and the corresponding threshold values for *k*^+^.

**Table 2.** Summary of the restrictions imposed by intrinsic *k*^+^ values on the values of the effective *on* and *off* kinetic rates of BCR-Ag (2D interactions) and Ab-Ag (3D interactions).

| Effective rates | Ranges of intrinsic $k^+$ | | |
| --- | --- | --- | --- |
| | $k^+ < 5 \times 10^6$ | $5 \times 10^6 < k^+ < 5 \times 10^8$ | $5 \times 10^8 < k^+$ |
| $k_{\text{eff}}^+(2D)$ | $\approx k^+$ | $\approx 10^6$ | $\approx 10^6$ |
| $k_{\text{eff}}^-(2D)$ | $\approx k^-$ | $[\frac{k_{\text{eff}}^-(3D)}{500}, \frac{k_{\text{eff}}^-(3D)}{5}]$ | $< \frac{k_{\text{eff}}^-(3D)}{500}$ |
| $k_{\text{eff}}^+(3D)$ | $\approx \frac{k^+}{5}$ | $\approx \frac{k^+}{5}$ | $\approx 10^8$ |
| $k_{\text{eff}}^-(3D)$ | $\approx k^-$ | $\approx k^-$ | $\approx \frac{10^8}{K_{A\text{eff}}(3D)}^a$ |

<sup>a</sup> $K_{A\text{eff}}(3D)$ , effective 3D affinity constant.

Contrary, the flexibility in the azimuthal angle *β* (Fig. 1c) is not well characterized. Nevertheless, based on the study of Zhang *et al* (20) that analyzed the conformational flexibility of IgG1 antibody molecules using individual particle electron tomography, one can argue that each paratope can cover a 180° range in the azimuthal angle. Thus, the total domain covered by both paratopes is Ω = {(*θ*, *β*)|*θ* ∈ [*θ*_min_, *θ*_max_], *β* ∈ [0, 2*π*]}. Furthermore, since *D*_*t*_(2*D*) ≫ *D*_*w*_(2*D*), we can neglect the marginal effect of the whole rotation motion, i.e 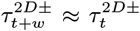. In this approximation, each paratope covers one half of Ω, namely Ω_1_ = {(*θ*, *β*)|*θ* ∈ [*θ*_min_, *θ*_max_], *β* ∈ [0, *π*]} and Ω_2_ = {(*θ*, *β*)|*θ* ∈ [*θ*_min_, *θ*_max_], *β* ∈ [*π*, 2*π*]}. For simplicity, we will suppose that the epitope is completely contained in either Ω_1_ or Ω_2_ and thus we only need to consider one of both paratopes (see Fig. S5).

Under the above definition of the domain Ω, the MFPTs depend on the initial position of the paratope (*θ*, *β*), and the epitope location *θ*_epi_ and size (*δ*; see Fig. 1c,e). Numerically, we model the rotational tilt diffusion combined with the whole rotation of the Fc region as a Brownian motion of the paratope on a spherical surface, which terminates when it reaches the epitope. Similarly, the rotational alignment step can be represented as a Brownian motion of the paratope in a circumference that terminates when it acquires an adequate orientation, that is, an angle orientation that allows binding. The mean *on* and *off* rotational times corresponding to the different rotational rate constants can be estimated using the theory of stochastic narrow escape (10) and first-passage processes (11), and are here denoted rotational mean first passage times (MFPT).

Through Monte-Carlo simulations, we computed the global MFPT 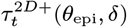 for the paratopes of the different IgGs subclasses, and for multiple values of *δ* = {19°, 14°, 10°, 6°, 2°} and different positions of *θ*_epi_. The results, summarized in Fig. 2, revealed conspicuous differences in the global MFPTs of the IgGs subclasses, being more notorious between the IgG2b and IgG1 subclasses. Interestingly, the difference between IgGs subclasses increases as the size of the paratope *5* decreases. In agreement with previous works (21), these results suggest that the same paratope in different Ab classes might result in different rotation rate constants and, therefore, in different effective rate constants, besides any differences imposed by Ig class-specific glycosylation (12, 22).

**Fig. 2.**
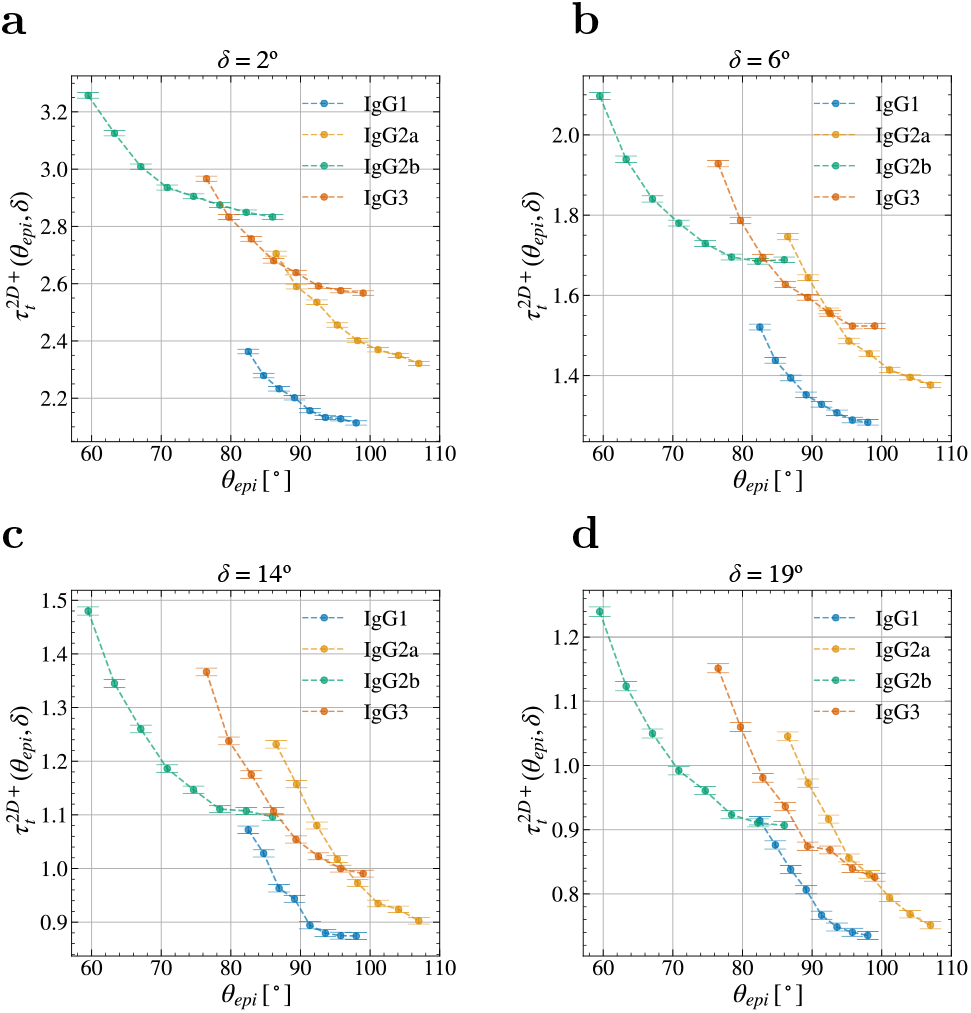
Isotype Fab tilt flexibility affects the effective orientational rates. MFPT 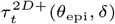 computed through Monte-Carlo simulations for the different mouse IgG subclasses and multiple values of *θ*_epi_ and *δ*. Parameters: *dt* = 2 · 10^−4^, *D* =1, # initial positions = 3000, # repetitions per initial position = 100.

The exact solution of the system of Eqs. (S1) for the MFPT 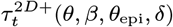 requires the knowledge of the pseudo-Green functions for the domains in Fig. S5 that, to our knowledge cannot be obtained using closed mathematical expressions. We use a variational method described in the Supplementary Information to overcome this limitation. The resulting *on* rate is given by

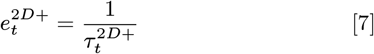

with 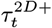 given by Eq. (S19).

The off-rate process for BCRs is similar to that of soluble Abs, except that the whole rotational motion marginally contributes to the diffusion of the paratope. Therefore, the rotation off-rate 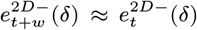 is given by Eqn. (S16) but with *D_t_* instead of *D*_*t*+*w*_:

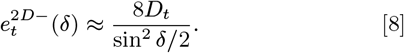

### Estimates for the rate constants 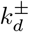 and *e*^±^ in 2D and 3D, and the corresponding *k*^+^ threshold values

Equations 2 and 3 allows connecting the underlying or intrinsic kinetic rates with the effective 2D and 3D ones. Thus, here we integrate all the computations in the preceding sections. Calculations for 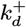 and 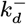 can be found in (7). However, to estimate the rate constants *e*^±^ it is necessary to know, in addition to the angles *ϕ*_0_ and *δ*, the rotational diffusion constants *D_a_*, *D_t_* and *D_w_* in 2D and 3D conditions. These diffusion constants can be calculated as follows. The mean square angular deviation of a rotating molecule after a time *t* is given by (19, 23)

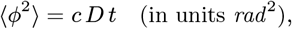

where *c* is a constant particular for the geometry of the rotation, and *D* is the corresponding rotational diffusion constant. In particular, *c* = 2 for Fab twist and BCR Fc axial rotations, and *c* = 6 for Fab tilt and whole Ab rotations (7, 24). For such rotations, the mean rotational correlation time, denoted *t*irad, is the average time spent by the molecule to rotate 1 rad. Therefore,

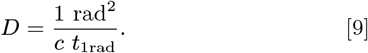

The values for *t*_1rad_ corresponding to the three rotations considered here (whole molecule, tilt and alignment) and/or the respective rotational diffusion constants, *D_w_*, *D_t_* and *D_a_*, estimated for Igs both in solution (3D) and attached to a large particle (2D) are summarized in Table S3. In addition, and as previously mentioned, although *D_w_* (2D) for the rotation of BCRs about the Fc axis has not been determined, one can take as a reference value that of the epidermal growth factor receptor (19), *D_w_*(2D) ≈ 1.43 × 10^3^ rad^2^/s.

We can now go beyond qualitative arguments and calculate the global rotational kinetic rates, *e*^+^ and *e*^−^. In particular, applying the estimations for the rotational diffusion constants in Table S3 and for the angles *φ*_0_ and *δ* (SI, section B) to Eqs. (S6, S12, S16, S19), and then applying the resulting values to Eq. (6). Ranges for the translational and global rotational kinetic rates in 2D and 3D conditions thus estimated, together with the corresponding threshold values for *k*^+^ given by Eq. (4), are summarized in Table 1.

These results show that at variance with 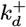 and 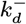, that have very different values from 2D to 3D conditions but similar scaling factors (~ 10^3^-fold), *e*^+^ and *e*^−^ have similar values in 2D and 3D conditions, with *e*^−^ being 2-fold lower in 2D than in 3D. However, what is most relevant here is that the *k*^+^ threshold values differ by ~ 10^3^-fold from 2D to 3D conditions. Hence, 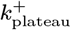 for 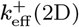 rates (BCR-Ag interactions) is much lower than for 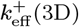 rates (free Ab-Ag interactions). More specifically, the 2D 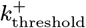 is ⩽ 10^6^ s^−1^ and for intrinsic binding rates *k*^+^ larger than this threshold, 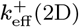 rate approaches a plateau of (1-2) ×10^6^ M^−1^ s^−1^. Hence, intrinsic *k*^+^ rates higher than that threshold cannot have a differential impact on the effective binding rate of Ag to B cells. In contrast, the 3D 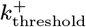 is ≈ 5 × 10^8^ s^−1^, and for values of the intrinsic binding rate *k*^+^ larger than this threshold the 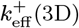 rate plateaus between 5 × 10^8^ and 5 × 10^9^ M^−1^ s^−1^ (left panel in Fig. 3).

**Fig. 3.**
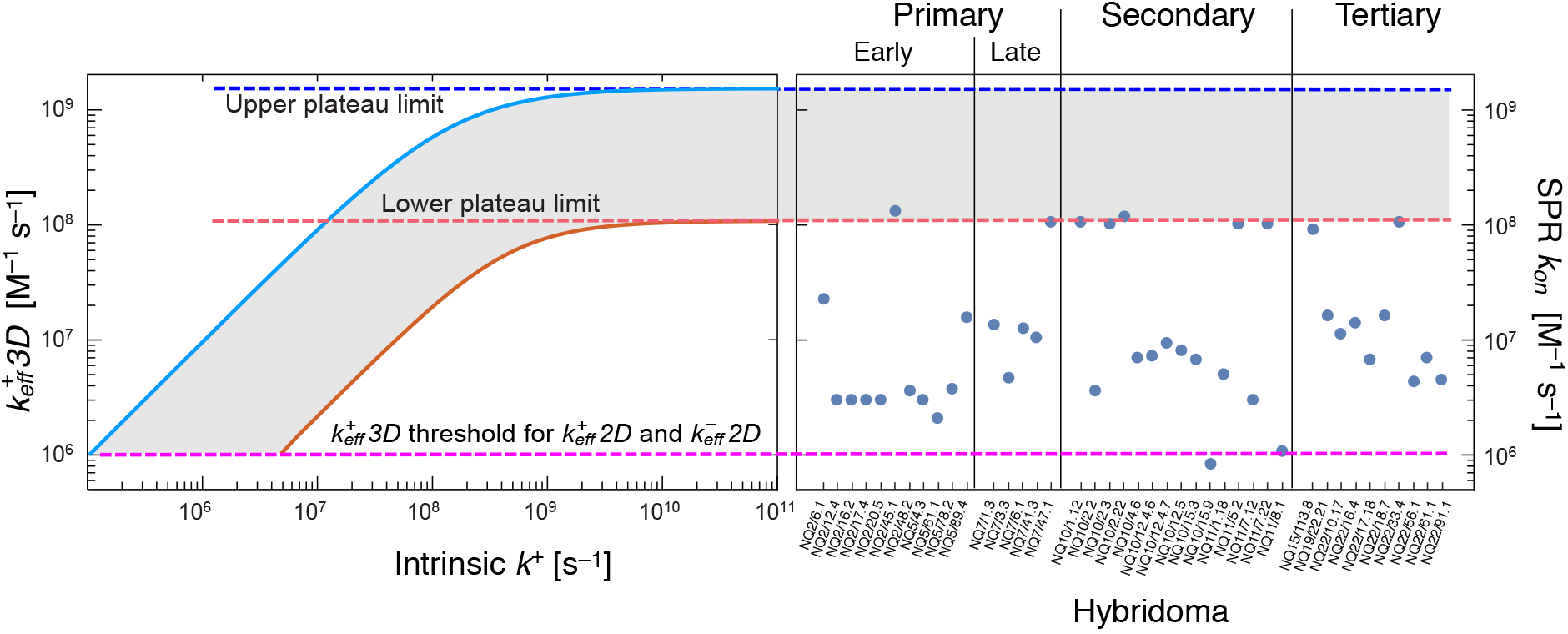
Translational and rotational diffusion events put a limit to effective 3D *on*-rates. Left panel, theoretical estimation of 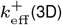 for Ab-Ag interactions as a function of the intrinsic binding rate, *k*^+^, based on Eq. (2). Dashed blue and red lines correspond, respectively, to upper and lower plateau boundaries. They are the mean ±3sd of the distribution of plateau values obtained with 10^4^ random sets of parameter values within the confident limits of the 3D translational and rotational rates in Table 1. Blue and red curves were obtained with the parameters set used to estimate, respectively, the lower and upper plateaus. The pink dashed line corresponds to the 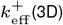 threshold for 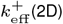 and 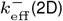 depicted in Fig. 4. Right panel, experimental 3D *k_on_* values obtained with the SPR assay for different anti-2-PhOx monoclonal Abs (redrawn from reference (25)). Note the close agreement between the highest experimental values and our predicted 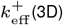 lower plateau. Notice that all experimental values, from an early primary immune response to a tertiary immune response, lay within the 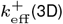 lower plateau and the 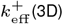 threshold.

## Upper limits to the effective kinetic rates of Ab- and BCR-Ag interactions

The above estimations greatly impact our understanding of the effective kinetics of BCR-Ag and Ab-Ag interactions and their mutual relationship.

It has long been made the empirical observation of a ceiling for the 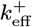 of monoclonal Abs and their ligands (see for instance (25, 26)). Calculations of the range for the 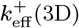 plateau of Ab-Ag interactions help to explain it. This is illustrated in Fig. 3 left side, where the mean variation of 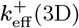 ±3 sd (solid lines) for increasing values of the intrinsic *k*^+^ rate and the range of the corresponding plateaus (dashed red and blue lines) are plotted. In this figure, we also plot side by side with experimentally observed 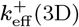 values for samples of monoclonal Abs obtained in primary, secondary, and tertiary murine immune responses against the hapten 2-phenyl 5-oxazolone (PhOx; Fig. 3 right side, based on (25)).

Given that 2D 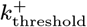 and 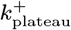 are at least 2 orders of magnitude lower than the corresponding 3D values, the 3D Ab-Ag effective affinity and rate constants cannot be used in general as surrogates for 2D BCR-Ag effective affinity and rate constants. However, this does not mean they cannot be used in all cases. For instance, for values of intrinsc 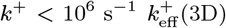 and 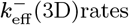rates are close to the expected 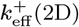 and 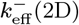 rates, respectively. This can be shown to be the case by expressing 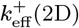 and 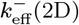 as a function of 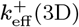 (see SI section E).

Plotting the ratios 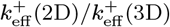 and 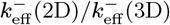 as a function of 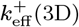 (Fig. 4) shows that 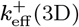 is a moderately reasonable proxy of 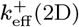 only for values of 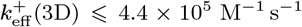. At that value 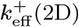 is in the range of 59% to 80% of 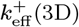. However, 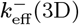 is a reasonable proxy of 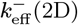 for values of 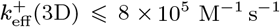 (at which value 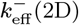 is in the range of 57% to 80% of 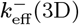). These are very important results as they establish the approximate limits up to which the 3D kinetic rates of Ab-Ag binding provide values close to the expected BCR-Ag 2D kinetic rates.

**Fig. 4.**
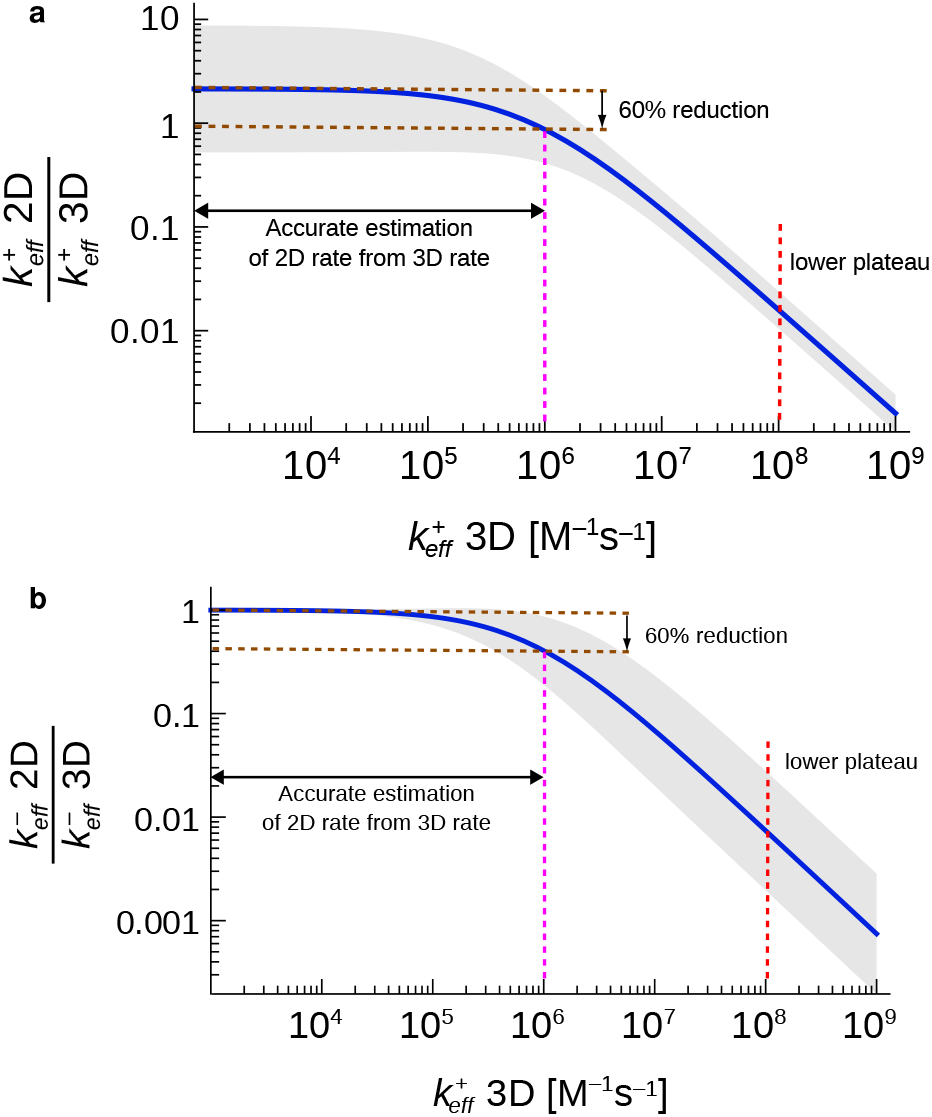
**3D Effective *on*-rate *observability thresholds* for estimating 2D effective *on* and *off* rates** using Eq. (S21) and Eq. (S22). Blue solid lines, variation of the ratios 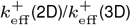 (**a**) and 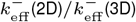 (**b**) as a function of 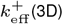. Each of these curves is an average from a distribution of curves obtained with the same 10^4^ random sets of parameter values used in Fig. 3. Grey areas correspond to curves within the mean ±2sd. A 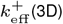 threshold for each curve was defined as that 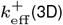 value for which a 60% reduction is obtained. This threshold is 10^6^ M^−1^s^−1^ for both ratios and it is indicated with a pink dashed line. For comparison, a red dashed line corresponding to the 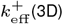 lower plateau in Fig. 3 is also depicted.

## Discussion and Conclusions

During B cell-FDC interactions, BCRs bind cognate Ag tethered on FDCs’ membrane (27) and hence in 2D conditions. In contrast, in *in vitro* assays, Ab-Ag interactions take place with at least one component in solution (e.g., SPR assays) and thus in 3D conditions. In the case of T lymphocytes, the consequences of these two biophysical conditions on receptorligand interactions have been analyzed to some extent. Experiments (28, 29) and mathematical models (6) show that the effective kinetic rates of TCR-ligand interactions obtained in 3D conditions cannot predict the effective kinetic rates obtained in 2D conditions (28, 29). Moreover, it was also shown that it is the 2D kinetics, not the 3D kinetics, that correlate with Ag potency to activate T cells (30, 31). The theoretical analysis of TCR-ligand interactions was general enough to apply to the cases of BCR-Ag and Ab-Ag interactions (7), but until now it was not possible to derive quantitative implications for the relationship between 2D BCR-Ag and 3D Ab-Ag interactions. This was due to a lack of quantitative biophysical data on the different modes of rotational diffusion of paratopes in BCRs and Abs. Here we have extended our previous work by performing a detailed theoretical analysis of available data on whole molecule, Fab tilt and Fab twist rotations, which allowed us to obtain, for the first time, quantitative estimates of the global *on* and *off* rotational rates of BCR and Ab paratopes.

To determine the parametric dependence of the different *on* and *off* paratope rotational rates on the structural properties of both BCRs and Abs we have used the Narrow Escape Theory (10) and First Passage Processes (11). We have characterized every step entailing the rotational degrees of freedom, so we could use the most precise available empirical estimates of Fab-Fab and Fab-Fc angles, and of mean rotational correlation times corresponding to the three rotations considered here. This has been a challenging task as empirical data on angle ranges and mean rotational correlation times for different Ig isotypes are scarce and scattered from a few mammal species, so we expect new targeted studies will improve and narrow our estimates.

Having derived quantitative values for the translational and rotational rates of paratopes enabled us to calculate the thresholds of the intrinsic *k*^+^ rate, in 2D and 3D conditions, beyond which 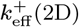 and 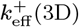 approach fast a plateau, as well as the actual values of these plateaus. In particular, the plateau for 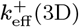 is fast approached for values of the intrinsic molecular *on* rate *k*^+^ > 10^9^ s^−1^ (Fig. 3, left panel). Although currently there is no experimental data of BCR-Ag interactions with which to compare the theoretical plateau for 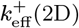, we could put to test the theoretical plateau we obtained for 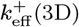 by comparing it with a rather comprehensive sample of monoclonal Abs anti-PhOx provided in (25). The lower and upper uncertainty limits of the theoretical 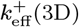 plateau are, respectively, ~ 10^8^ and ~ 10^9^ M^−1^s^−1^ (respectively, dashed red and blue lines in Fig. 3). Strikingly, the lower plateau limit of the theoretical 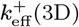 uncertainty band (grey area in Fig. 3) is in quite good agreement with the highest 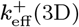 empirical values of the monoclonal Abs obtained from primary to tertiary immune responses to PhOx (Fig. 3, right panel), which lends strong support to our approach. Nevertheless, a wider sample of different sets of monoclonal Abs against different Ags will determine the generality or limitations of our approach. Yet, perhaps more surprising, and with deeper practical implications discussed below, is that the expected effective *on* and *off* rate ratios 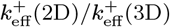 and 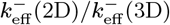 can be estimated as a function of 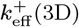 (Fig. 4). This is highly relevant because the 2D effective rate constants are the actual ones that determine B cell behavior. Thus, we have uncovered that for values 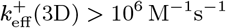, both 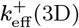 and 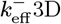 become bad estimators of the corresponding 2D effective constant rates (Fig. 4, vertical pink, dashed lines).

It is somewhat surprising that despite the widespread use of the seminal works by Berg and Purcell (14) and De Lisi (13), there has been put so little emphasis on the understanding of the biophysics of the molecular interactions of the two central actors in the humoral immune response, BCRs and Abs, with their cognate Ags. Our present calculations aim to stimulate experimental studies to establish the validity of the uncovered differences between effective BCR-Ag and Ab-Ag binding kinetics and to encourage the development of state-of-the-art molecular dynamics simulations and experimental methods to narrow the estimated rates for some of the aforementioned rotational events.

In summary, we have shown here that, contrary to the traditional implicit assumption, the kinetics of BCR-Ag interactions cannot be inferred from those of Ab-Ag interactions but for those with 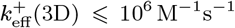. This has important implications in interpreting observations related to B-cell selection in germinal centers, an essential process in T-dependent humoral responses based on the amount of Ag endocytosed by B cells and presented to Tfh cells. For instance, above the thresholds established by our calculations for the 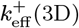 rate (Fig. 4, vertical pink, dashed lines), they become little informative —if at all— with respect to the Ag binding strength sensed by a germinal center B cell, and hence on the amount of Ag it can endocytose. In other words, for Abs with 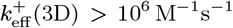 it cannot be inferred that the germinal center B cells that produced those Abs were selected because they had a higher 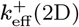 rate (or a lower 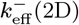 rate) than B cells producing Abs with 10-fold lower 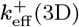 rates (or 10-fold higher 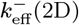 rates). We must be aware that, paraphrasing Malissen and Bongrand (32), a B cell can make decisions based only on parameters it senses, not on quantities measured *in vitro* by experimentalists. In this respect, the present results only concern one of two aspects usually neglected of the interpretation of the Ab affinity maturation process and the selective process(es) that lead to it. The other neglected aspect is the role of naturally generated tensile forces acting on Ag-bound BCRs, which can alter to a significant extent the intrinsic unbinding rate *k*^−^, similar to what has been shown for TCRs (33). This second aspect of the BCR-Ag interactions also deserves to be experimentally and theoretically studied.

## ACKNOWLEDGMENTS

This work has been partially funded by MCIN/AEI/ 10.13039/501100011033, through Grants PID2019-106339GB-I00, and PRE2020-092274, and by ESF Investing in your future, and by Xunta de Galiza under project GRC-ED431C 2020/02.

## Supporting Information

### Supporting Information Text

#### A. Flexibility of Fab arms for different Ig isotypes

Human IgG subclasses can be ordered in decreasing flexibility as: IgG3 *>* IgG1 *>* IgG4 *>* IgG2 (1). In total agreement with the reported functional equivalence between mouse and human IgGs (see Table S1) murine IgG subclasses can be similarly ordered as: IgG2b *>* IgG2a *>* IgG1 *>* IgG3 (2).

**Table S1.**
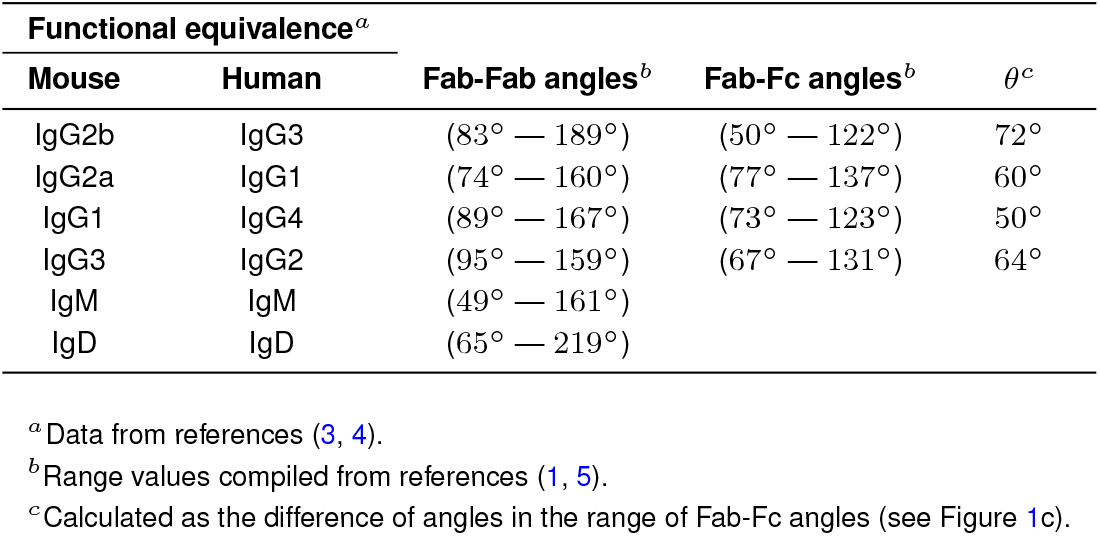
Empirical estimates of the tilt angle ranges of murine and human immunoglobulins. Note the agreement between the reported Fab-Fc tilt *θ* angles^*c*^ of mouse and human IgG subclasses.

#### B. Mathematical derivation of the rotational rates

According to the theory of stochastic narrow escape (6) and first-passage processes (7), the mean first passage time (MFPT) until absorption of a Brownian particle inside a domain Ω, with absorbing *∂*Ω_*a*_ and reflecting *∂*Ω_*r*_ boundaries satisfies the following mixed boundary value problem:

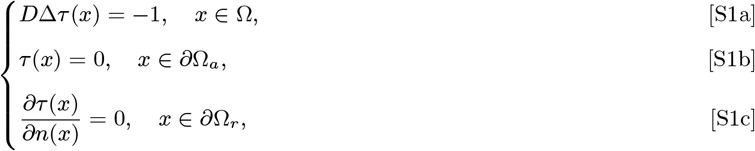

where *D* is the diffusion coefficient and **n** is the unit outer normal vector to the boundary. The MFPT *τ*(*x*) is dependent on the initial position *x* and the size of the absorbing domain Ω_*a*_, which in our case is determined, for the tilting step, by the angle *δ* corresponding to the epitope length, and, for the alignment step, by the angle range of adequate orientations *φ*_0_. From now on, we will be referring to *τ*(*x*, *δ*) or *τ*(*x*, *φ*_0_) as the MFPT and *τ*(*δ*) as the global MFPT. The inverse of *τ*(*δ*) is the desired on/off-rate and results from averaging over all possible initial positions. We summarize the domains and notations associated with each process in Table S2 where we also include a column pointing to the analytical solution of each sub-problem.

The angles *δ* and *ϕ*_0_ are those defined in Figures 1c and 1d in the main text. The tolerance deviation angle, *ϕ*_0_, has been estimated to be in the range 5°(≡ 0.087 rad) < *ϕ*_0_ < 11°(≡ 0.192 rad) (8). To estimate *δ*, we consider the spherical surface with radius *R* ≈ 84Å defined by a Fab arm (9, 10). On this surface, an epitope with length *L* = 28Å (11) defines a maximum angle *δ* ≈ 19°(≡ 0.332 rad).

**Table S2.** Description and geometry of the domains (Ω), absorbing domains (Ω_*a*_) and reflecting boundaries (*∂*Ω_*r*_) for binding and unbinding processes in B-cell receptors and free antibodies.

| Step<sup>a</sup> | Notation<sup>a</sup> | $\Omega$ | $\partial\Omega_a$ $\Omega_a$ | $\partial\Omega_r$ | Eqn. |
| --- | --- | --- | --- | --- | --- |
| Fab Alignment (+) | $e_a^+(\phi_0)$ | Arc $(\phi_0, 2\pi)$ | Arc $[0, \phi_0]$ | — | Eq. (S5) |
| Fab Alignment (—) | $e_a^-(\phi_0)$ | Arc $(0, \phi_0)$ | Arc $[\phi_0, 2\pi]$ | — | Eq. (S6) |
| Ab+Fab rotation (+) | $e_{t+w}^{3D+}(\delta)$ | Spherical cap $(\pi - \delta)$ | Spherical cap $(\delta)$ | — | Eq. (S12) |
| Ab+Fab rotation (—) | $e_{t+w}^{3D-}(\delta)$ | Spherical cap $(\delta)$ | Spherical cap $(\pi - \delta)$ | — | Eq. (S16) |
| BCR+Fab rotation (+) | $e_{t+w}^{2D+}(\delta)$ | Portion of Sphere | Spherical cap $(\delta)$ | $\theta \in \{\theta_{\min}, \theta_{\max}\}$<br>$\beta \in \{0, \pi\}$ | Eq. (7) |
| BCR+Fab rotation (—) | $e_{t+w}^{2D-}(\delta)$ | Spherical cap $(\delta)$ | Spherical cap $(\pi - \delta)$ | — | Eq. (8) |

<sup>a</sup> On and off rates are indicated, respectively, with symbols (+) and (—).

##### Alignment rotational 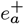 and 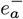

The *off* and *on* alignment rates, denoted here, respectively, 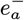 and 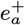, are in principle independent of the 2D and 3D conditions. For the off-rate process, the particle representing a twisting paratope diffuses within a range or arc of adequate orientations Ω^−^ = [0, *ϕ*_0_] until it loses a proper alignment through the boundaries 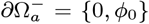. Whereas, in the on-rate process, the paratope diffuses inside an arc Ω^+^ = [0, 2*π* – *ϕ*_0_] until it acquires an adequate alignment through the boundaries 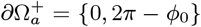. Denoting *ϕ*_−_ = *ϕ*_0_ and *ϕ*_+_ = 2*π* — *ϕ*_0_, the mixed boundary value problems for the associated MFPTs 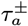 are:

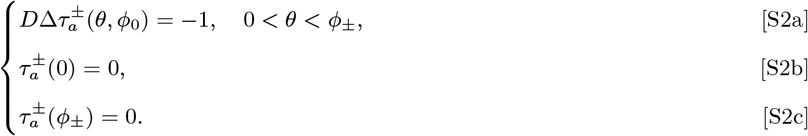

Problem (S2) can be solved through direct integration:

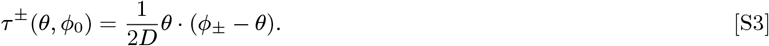

Averaging Eqn. Eq. (S3) over all initial positions *θ* ∈ [0, *ϕ*_±_] yields the global MFPTs:

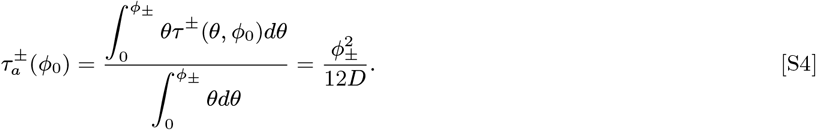

The corresponding *on* and *off* rates are given by the equations:

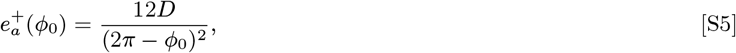

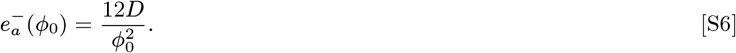

where *D* ≡ *D_a_*.

**Fig. S1.**
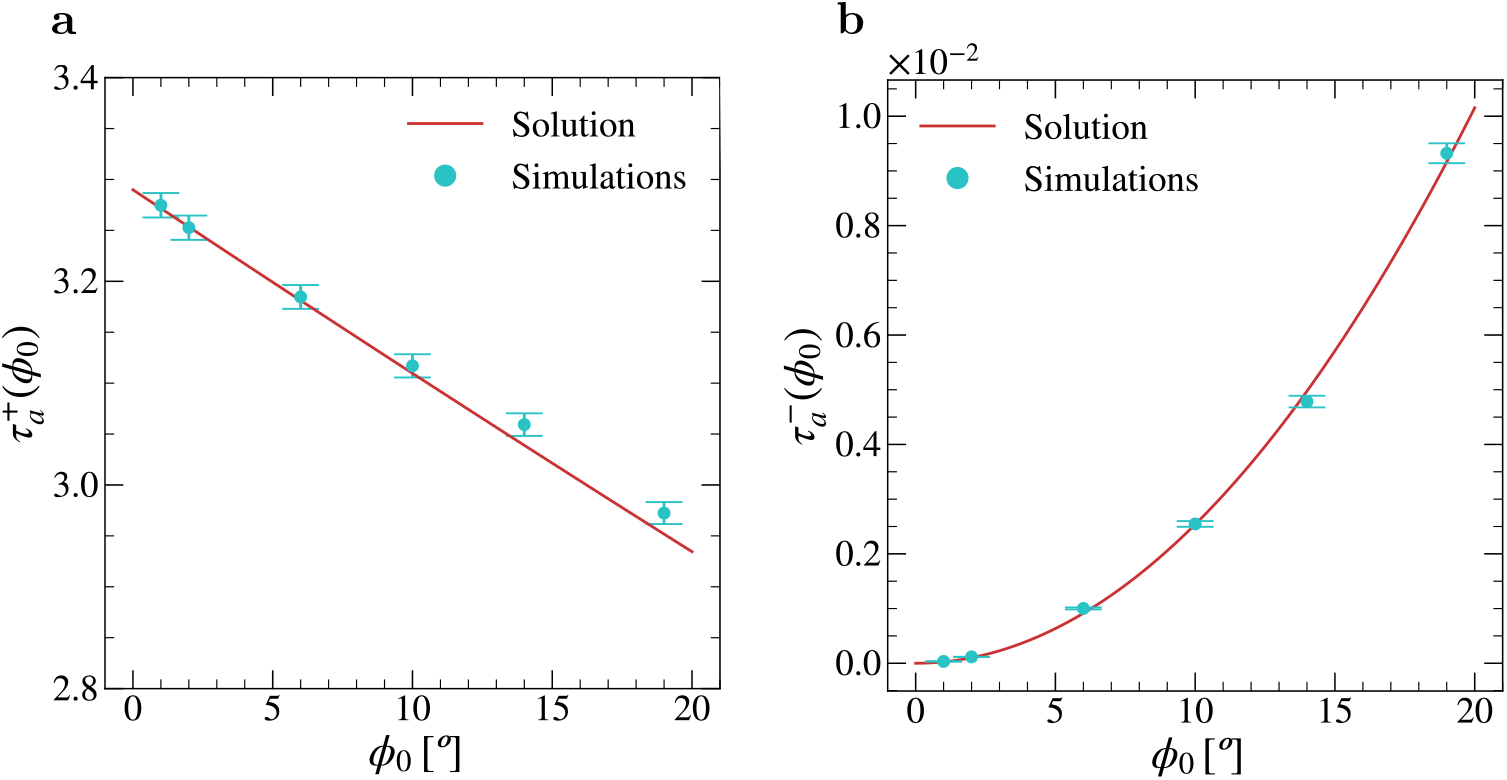
Comparison between explicit solutions for rotational alignment. (given by Eq. (S4)) and numerical calculations of the global MFPTs 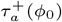 (**a**) and 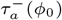 (**b**). Parameters in **a** and **b** for each value of *ϕ*_0_: *dt* = 10^−5^, *D* =1, # initial positions = 500, # repetitions per initial position = 2000. Error bars correspond to ±1sd.

Figure S1 shows a comparison between the times 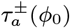 computed through Monte-Carlo simulations and the times obtained using the explicit solutions given by Eqn. (S4).

##### 3D rotational rates of Abs

For the on-rate 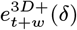, the particle representing the paratope diffuses throughout the spherical surface until it encounters the epitope represented by a spherical patch of angle *δ* (Fig. S2a). By setting the epitope at the north pole of the sphere, we can see that the MFPT until absorption is symmetrical in the azimuthal angle *β*, 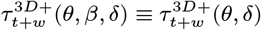. The mixed boundary value problem (S1) is, then, given by:

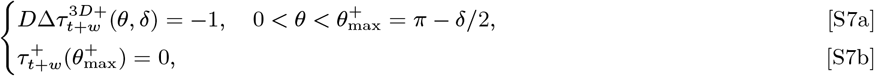

Note that for the off-rate, 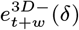, the particle diffuses inside a polar cap of angle *δ* (see Fig. S2b). Thus, the mixed boundary problem for 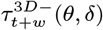 is equivalent to Eqn. (S7) but with the boundary set at 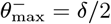, that is:

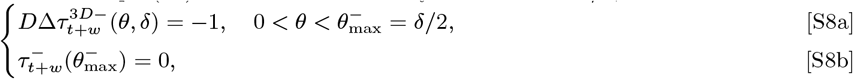

**Fig. S2.**
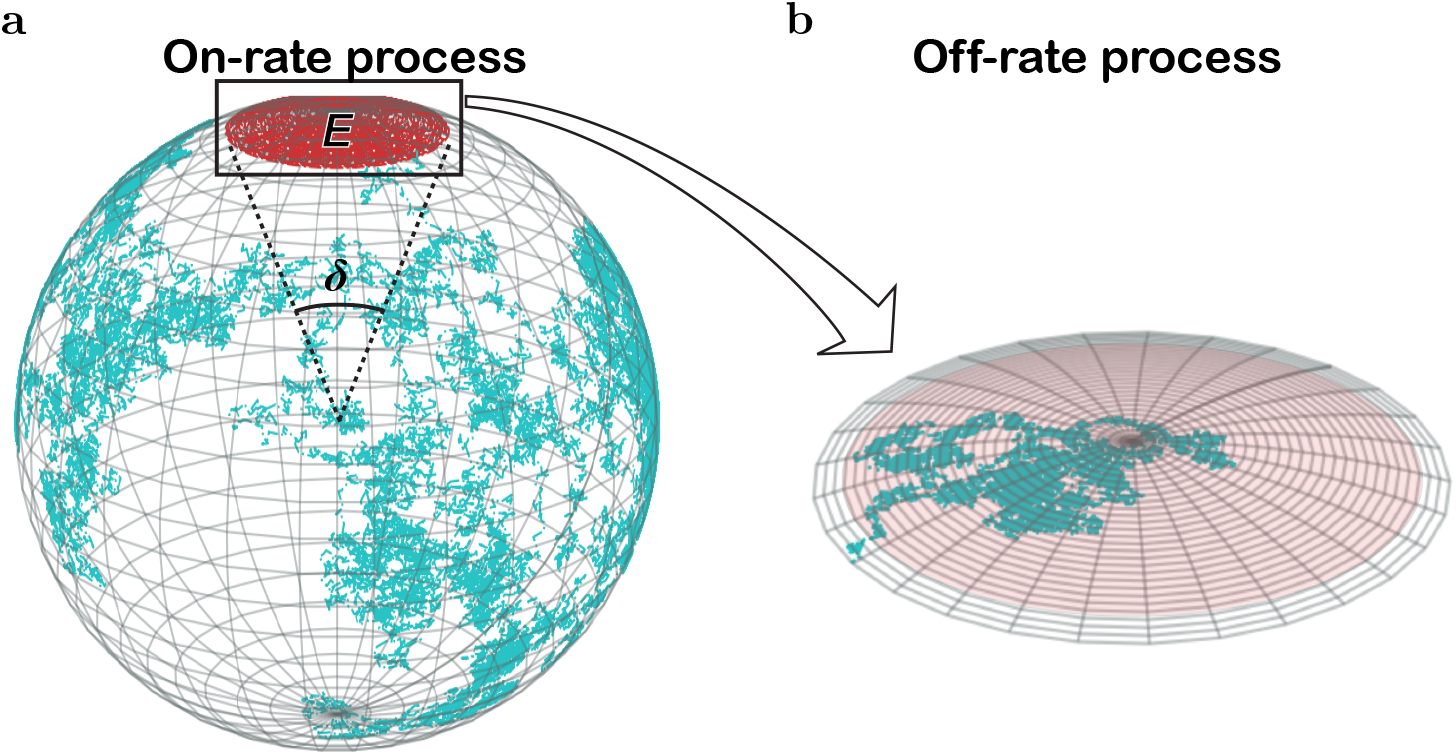
Random Walk view of a paratope diffusion. **a**, On-rate process: The center of mass of a paratope diffuses on the sphere (cyan broken line) until finding the epitope (red). *δ*, angle of the arc with length equal to the major length of the epitope. **b**, Off-rate process: The paratope center of mass diffuses within the epitope area (pale red) until it escapes (grey area).

Problems defined by Eqns. (S7–S8) can be solved by direct integration. For that, applying the change of variable *r* = sin *θ* and substituting the Laplace operator in spherical coordinates yields:

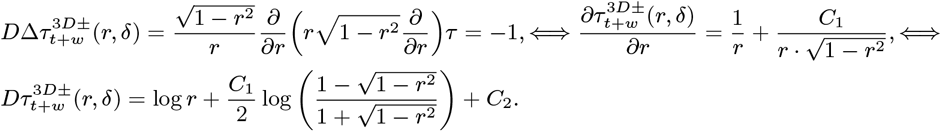

Substituting back *r* by sin *θ* gives the equation:

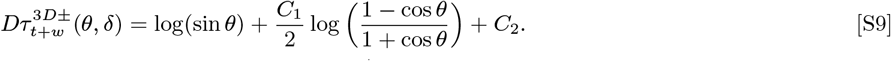

The value of *C*_2_ is obtained from the absorbing boundary condition at 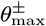, and thus:

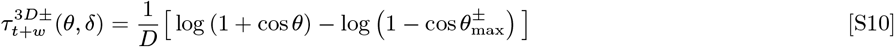

A comparison of the simulated MFPT 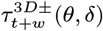 with its exact value given by Eqn. (S10) is shown in Fig. S3. Finally, averaging Eqn. (S10) over all possible initial positions *θ* ∈ [0, *θ*^±^] gives the following expression for the global MFPT:

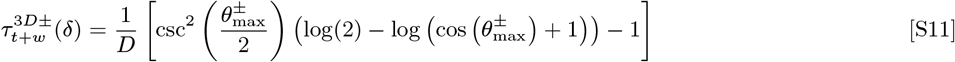

and, thus, the associated mean *on*-rate follows the equation:

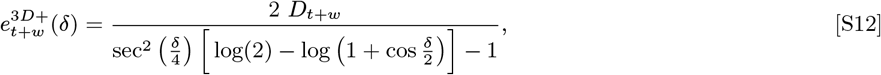

where *D* ≡ *D*_*t*+*w*_ ≈ *D_t_* + *D_w_*.

For the *off*-rate 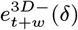, since the angle size of the epitope is relatively small (*δ* ≈ 19°), we can approximate the off-rate taking the limit *δ* → 0. In this limit, the epitope can be considered as a disk of radius *R* = sin *δ*/2, thus the problem for the mean first passage time 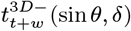 (sin *θ*, *δ*) is:

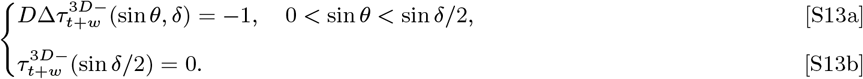

**Fig. S3.**
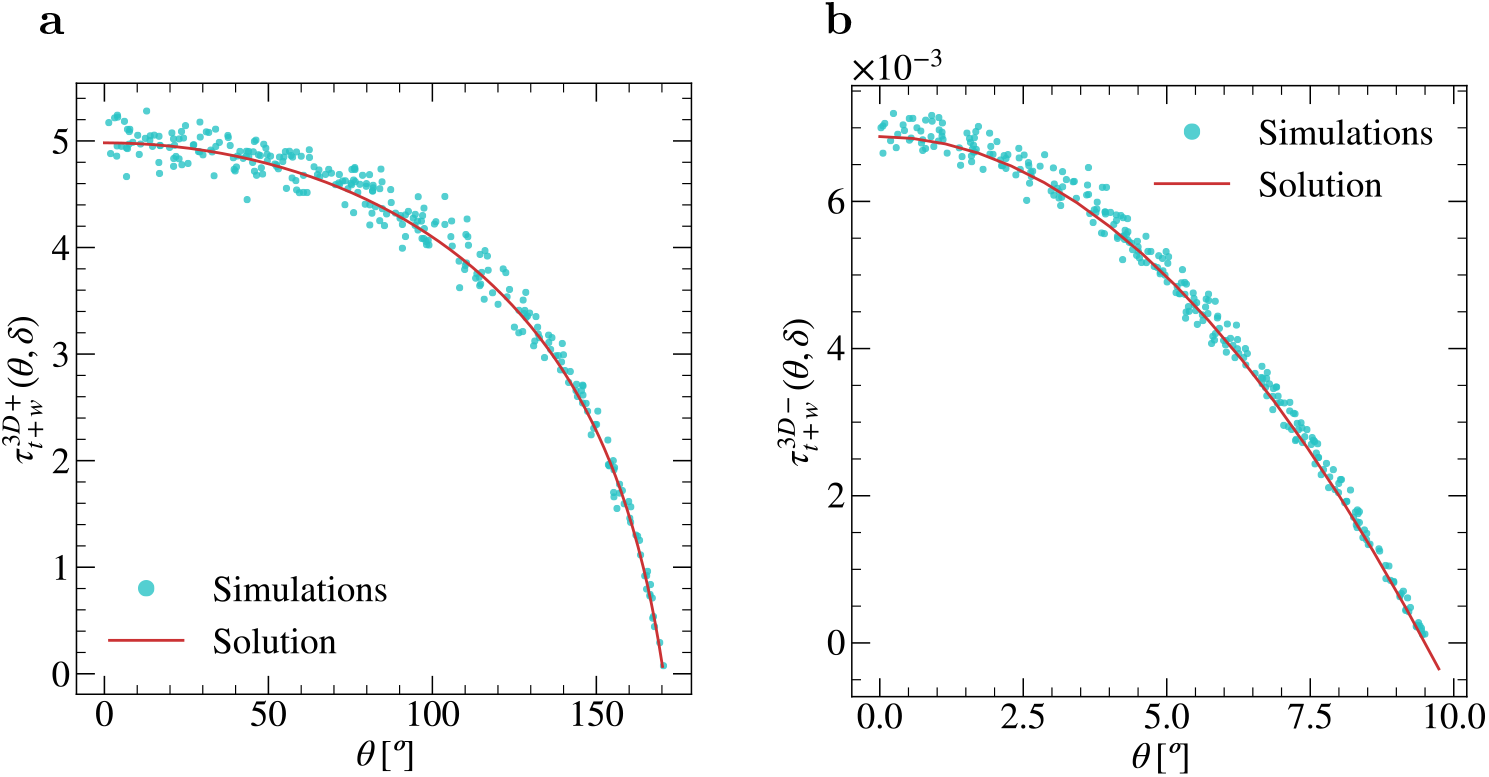
Comparison between MFPT numerical calculations and explicit solutions for 3D whole Ab and Fab tilt rotations. Simulated 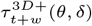 (**a**) and 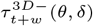 (**b**) (cyan dots) and their explicit solution (red lines) given by Eq. (S10). Parameters: **a**, *δ* = 19°, *dt* = 10^−5^; **b**, *δ* = 19°, *dt* = 10^−6^; # initial positions = 300; # simulations per initial position = 1000.

**Fig. S4.**
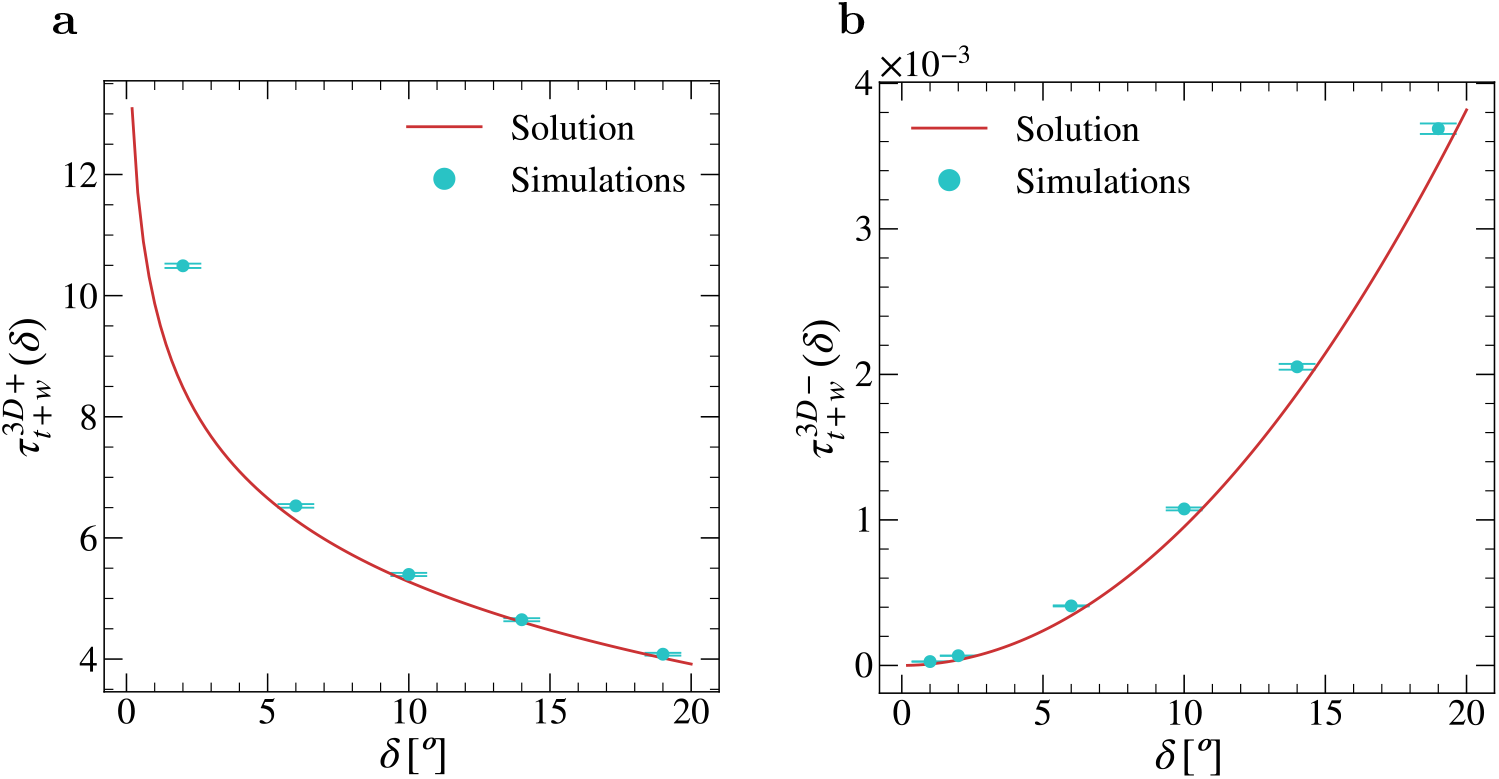
Comparison between global MFPT numerical calculations and explicit solutions for 3D whole Ab and Fab tilt rotations. Simulated (cyan dots) 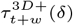 (**a**) and 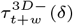 (**b**) and their explicit solutions (red lines) given by Eqs. (S11, S15). Parameters in **a** for each value of *δ*: *dt* = 2 × 10^−4^, *D* = 1, # initial positions = 3000, # repetitions per initial position = 100. Parameters in **b** for each value of *δ*: *dt* = 10^−5^, *D* = 1, # initial positions = 3000, # repetitions per initial position = 1000. Error bars correspond to ±1sd.

Applying the change of variable *r* = sin*θ*, Eqn. (S13) yields:

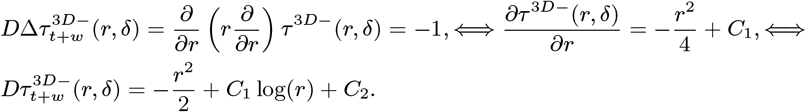

From the condition of finitude at *r* = 0, it follows that *C*_1_ = 0, and from the absorbing boundary condition it follows *C*_2_, which yields:

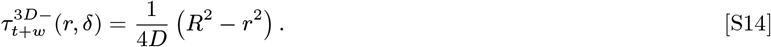

Averaging over the initial position *r* ∈ [0, *R*] gives the global MFPT:

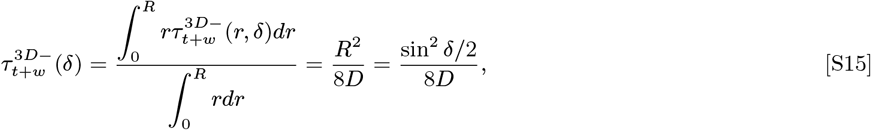

and, therefore, the corresponding approximated off-rate is:

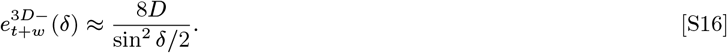

with *D* ≡ *D*_*t*+*m*_.

A comparison of the simulated global MFPT 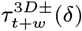 with its exact value given by Eqns. (S11, S15) is shown in Fig. S4.

**Fig. S5.**
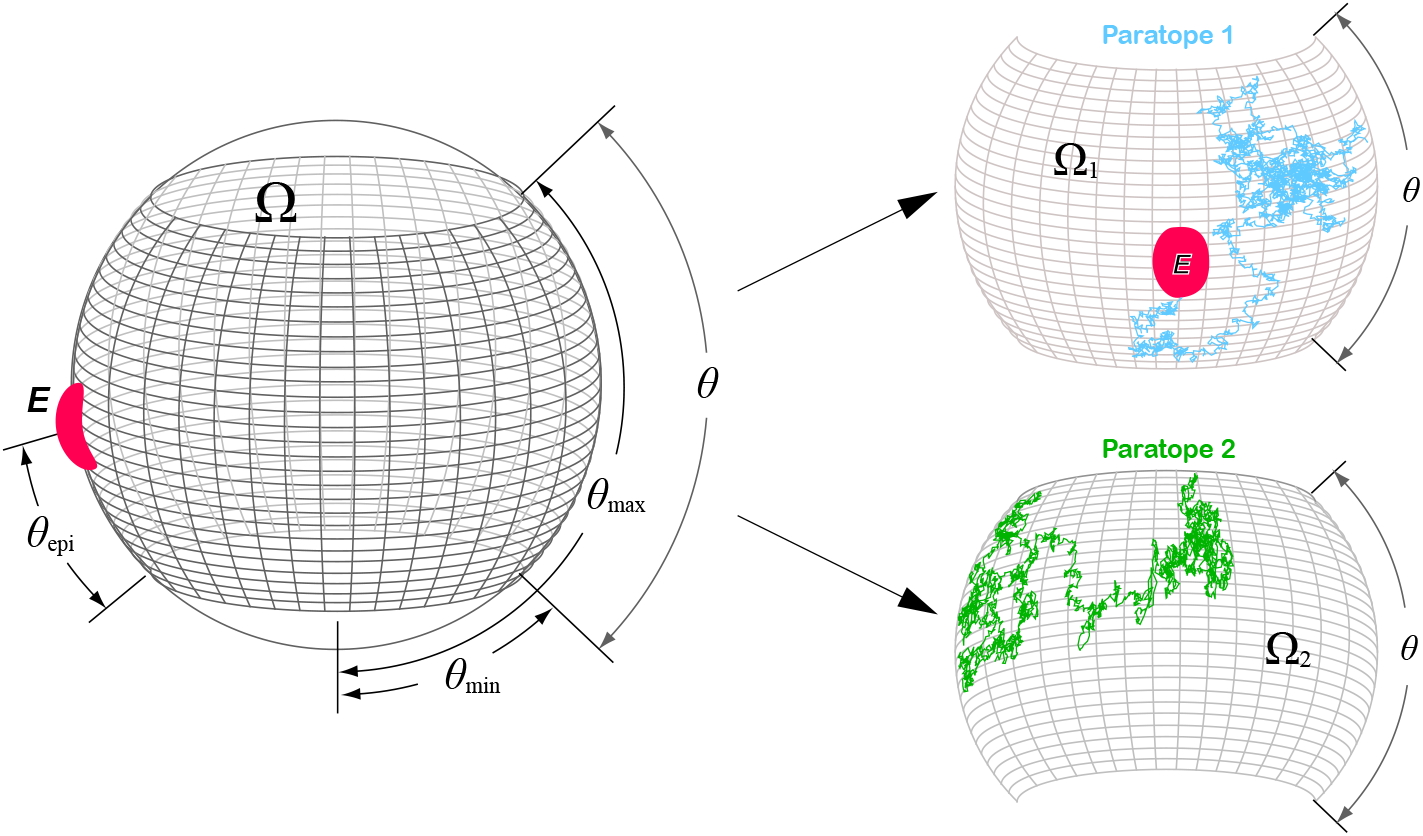
Spatial constraints for BCRs paratope tilt rotation. Neglecting the whole rotation motion, each paratope of a BCR covers only one half (Ω_1_ and Ω_2_) of the whole domain Ω = Ω_1_ ∪ Ω_2_. If the epitope (*E*, depicted in red) is completely contained in Ω_1_ or Ω_2_ then only one paratope contributes to the search for the epitope (cyan broken line in subdomain Ω_1_ and green broken line in subdomain Ω_2_).

#### C. Spatial constraints for BCRs paratope tilt rotation

##### Variational method

We will consider the case where the epitope is at the north pole of the sphere and the Fab tilt is only limited in southern regions and leave one free parameter to improve the accuracy of the approximation. In the latter situation, the MFPT only depends on the size of the epitope *δ* and the initial position *θ* ∈ [*δ*/2, Δ*θ*_eff_], where 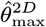 is the *variational angle* that should be related with the arc length 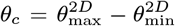 of the original problem. In Fig. S6a we show the remarkable agreement between the variational approximation and the numerical simulations. The mixed boundary value problem for the MFPT 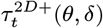 is defined by the set of equations (S1) but applied to 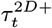:

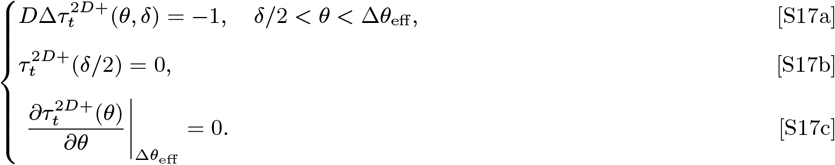

Problem (S17) has the same structure as problem (S7) with the addition of the reflecting boundary condition at Δ*θ*_eff_. Applying the boundary conditions, Eqns. (S11b–c) gives the solution:

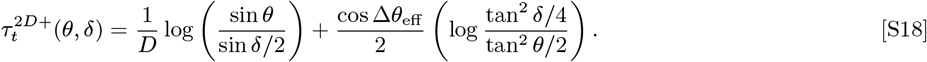

Averaging over the initial positions *θ* ∈ [*δ*/2, Δ*θ*_eff_] gives the global MFPT:

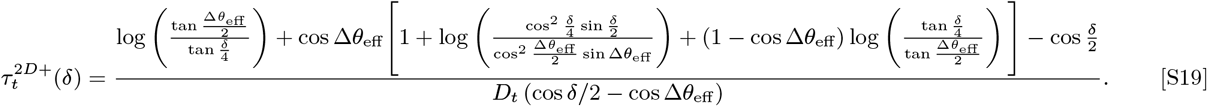

The resulting on rate is given by Eq. (7).

Figure S6b shows the estimated global MFPT 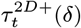 of the IgGs subclasses averaged over the different positions of *θ*_epi_ and a non-linear least square fitting to Eqn. (S19) with respect to the free parameter Δ*θ*_eff_. The fitted values of Δ*θ*_eff_ correlate linearly with the angle difference 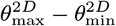 (*R*^2^ = 0.98, 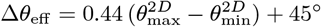).

**Fig. S6.**
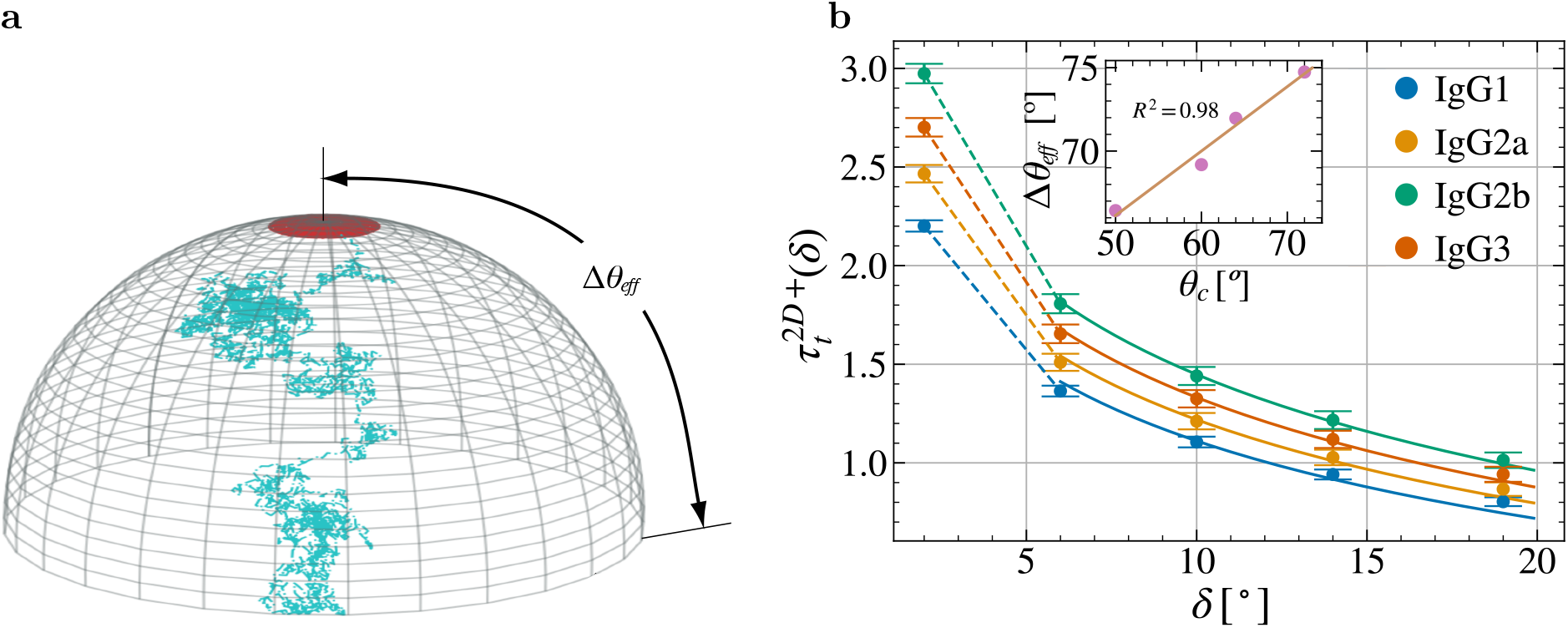
Variational Global MFPT approximation of 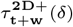. **a**, As an approximation of BCR’s paratope diffusion process (Fig. S5), we consider the case where the epitope is at one of the poles and the paratope can diffuse in a polar cap of size Δ*θ*_eff_. **b,** Global MFPT 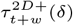 for the different IgGs subclasses. Single points are the computed global MFPT 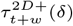, which result from averaging 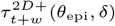 of Fig. 2 over *θ*_epi_. Solid lines are given by a non-linear least square fitting to Eqn. (S19). Inset shows the values of the fitted Δ*θ*_eff_ against the difference *θ_c_* = *θ*_max_ – *θ*_min_ of each IgGs subclasses. The point with *δ* = 2 was not considered for the fitting due to its unfeasibility in real epitopes. Values of Δ*θ*_eff_: 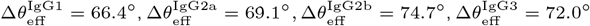. Error bars correspond to ±1sd.

#### D. Estimation of the 2D and 3D rotational diffusion constants, *D_w_*, *D_t_* and *D_a_*

The experimentally derived *t*_1rad_ values corresponding to the three rotations considered here (whole molecule, tilt and alignment), estimated for Igs both in solution) (3D) and attached to a large particle (2D), and the rotational diffusion constants, respectively, *D_w_*, *D_t_* and *D_a_*, theoretically derived using Eq. 9, are summarized in Table S3.

#### E. Estimation of 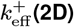 and 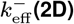 as a function of 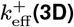

We first use Eq. (2) corresponding to the 3D case to express the intrinsic *k*^+^ in terms of the 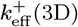 rate,

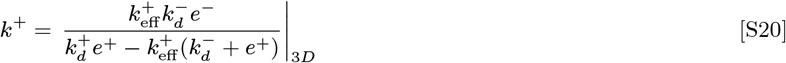

and then substituting this expression for *k*^+^ in the 2D case of Eq. (2), we get:

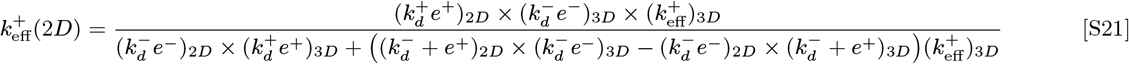

Similarly, using Eqs. (2) and (3) corresponding to the 3D case to express, respectively, *k*^+^ and *k*^−^ in terms of 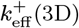 and 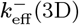 and then substituting the resulting expression for *k*^+^ and *k*^−^ in the 2D case of Eqn. (3), the following expression for 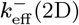 in terms of the effective rates 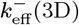 and 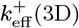 is obtained:

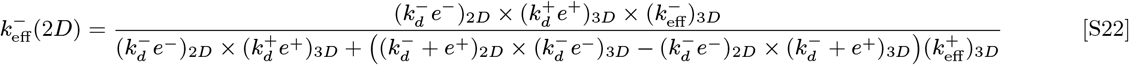

**Table S3.**
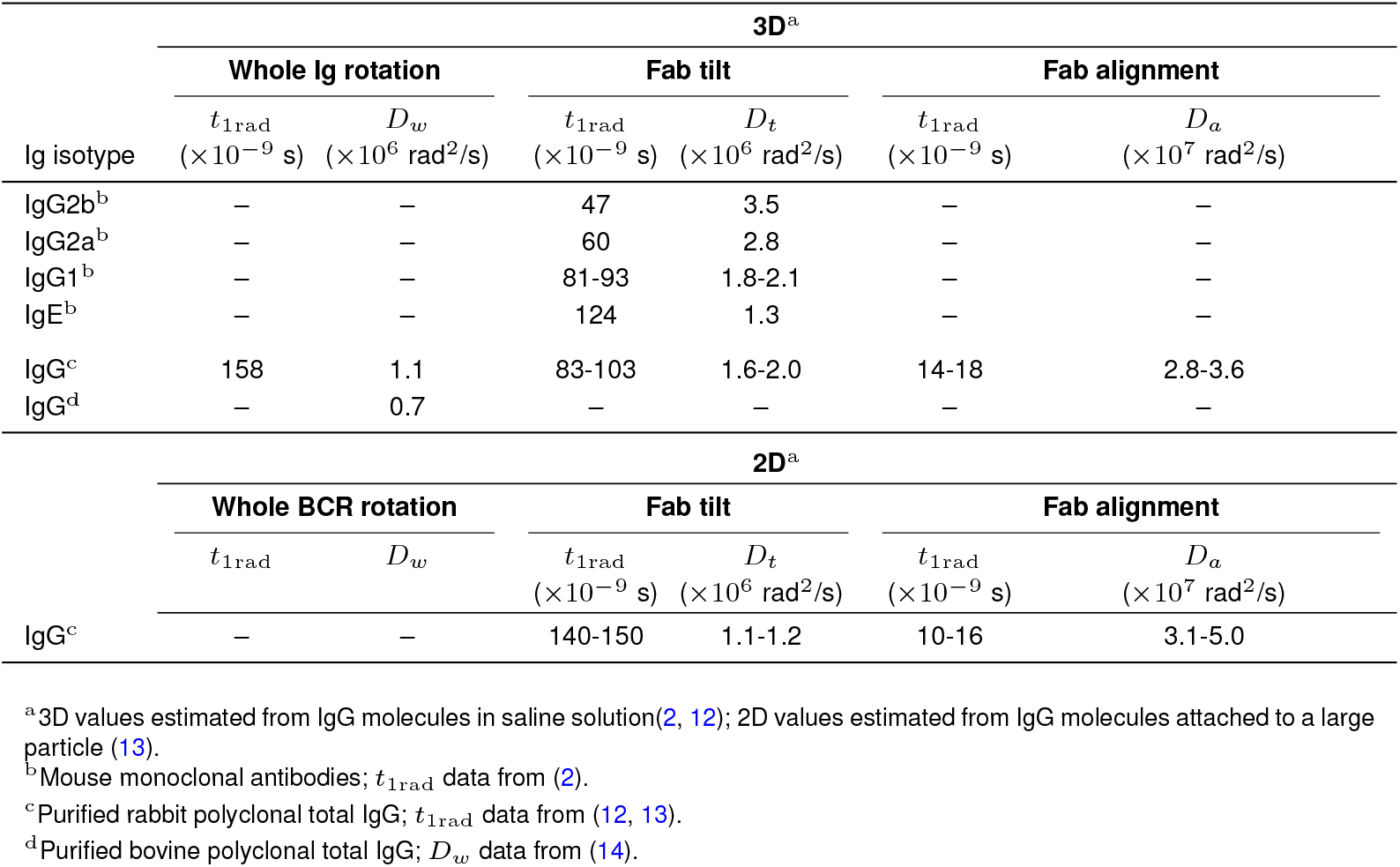
Mean rotational correlation times of IgG molecules and *D_t_* and *D_a_* diffusion constants.

